# Characterization of nitrogen deposition by means of atmospheric biomonitors

**DOI:** 10.1101/118257

**Authors:** E. A. Díaz-Álvarez, E. de la Barrera

## Abstract

An increase of nitrogen deposition resulting from human activities is not only a major threat for global biodiversity, but also for human health, especially in highly populated regions. It is thus important and in some instances legally mandated to monitor reactive nitrogen species in the atmosphere. The utilization of widely distributed biological species suitable for biomonitoring may be a good alternative. We assessed the suitability of an ensemble of atmospheric biomonitors of nitrogen deposition by means of an extensive sampling of a lichen, two mosses, and a bromeliad throughout the Valley of Mexico, whose population reaches 30 million, and subsequent measurements of nitrogen metabolism parameters. In all cases we found significant responses of nitrogen content, C:N ratio and the δ^15^N to season and site. In turn, the δ^15^N for the mosses responded linearly to the wet deposition. Also, the nitrogen content (R^2^ = 0.7), the C:N ratio (R^2^ = 0.6), and δ^15^N (R^2^ = 0.5) for the bromeliad had a linear response to NOx. However, the bromeliad was not found in sites with NOx concentrations exceeding 80 ppb, apparently of as a consequence of exceeding nitrogen. These biomonitors can be utilized in tandem to determine the status of atmospheric nitrogenous pollution in regions without monitoring networks for avoiding health problems for ecosystems and humans.

## Introduction

Nitrogen deposition is one the most predominant forms of atmospheric pollution^1^. This phenomenon results from the release of nitrogenous compounds to the atmosphere, both in cities, and in the country, including oxidized (NOx) and reduced (NHx) species, which are highly reactive. Nitrogen deposition causes acidification and eutrophication of aquatic and terrestrial ecosystems, contributes to the proliferation of invasive species leading to changes in ecosystem structure; it is also an important factor for climate change^1-6^. Nitrogenous pollution is also an issue for human health worldwide. Indeed, in traffic-jammed motorways, high accumulations of NOx at ground level are harmful by direct inhalation^7^. NOx emissions also contribute to the formation of secondary compounds including tropospheric ozone and particulate matter which are responsible in part for 9 million annual deaths worldwide^8^. For the case of Mexico City, a megalopolis with more than 20 million habitants, at least 9600 deaths can be attributed annually to atmospheric pollution, resulting from respiratory and heart diseases, and pollutant build up in the nervous system and the brain ^9,10^. In turn, ammonia emissions from agricultural sources quickly react with other atmospheric pollutants to form airbone particulate matter that can be readily inhaled^7^. Yet nitrogen is mostly neglected in public policy despite its importance in the prevalent discourse of global environmental change.

While monitoring nitrogenous pollution is an important issue, the deployment of air quality monitoring networks can be cost prohibitive in regions with developing economies. To fill this gap, an affordable alternative is the use of biomonitors^11^. For instance, throughfall deposition has been determined by the nitrogen content of lichen thalli^12^. Mosses are also widely used as biomonitors that allow covering vast areas, given that their nitrogen content reflects the rates of deposition and their isotopic composition can reflect the possible sources of pollution^13,14^. A group of special promise for biomonitoring in the neo-tropical region are atmospheric bromeliads, whose succulent tissues allow for yearlong physiologically activity, taking up pollution regardless of the seasonal environmental conditions. This is the case for the genus *Tillandsia* (Bromeliaceae), amply distributed in the Americas, which has been utilized for monitoring NOx pollution given that its nutrition depends exclusively from atmospheric sources^15,16^. However, physiological limitations of these atmospheric organisms can confuse the observed deposition, therefore it has been proposed that an assemble of various biomonitors for tracking this type of atmospheric pollution improves biomonitoring^11^.

To assess whether the combined use of various biomonitors can provide reliable information on nitrogen deposition we: 1) determined the spatial distribution of nitrogen content and the isotopic composition of the lichen *Anaptychia* sp., the mosses *Grimmia* sp., and *Fabronia* sp., and the bromeliad *Tillandsia recurvata* throughout the Valley of Mexico; and 2) evaluated the suitability as biomonitors of these organisms by comparing their nitrogen with an existing automated atmospheric monitoring network.

## Results

### Spatial distribution of nitrogen deposition

The total wet deposition in Mexico City increased from east to west, it ranged from 23 to 45 kg ha^−1^ year^−1^ during 2013, and between 24 and 50 kg ha^−1^ year^−1^ during 2014 (Figure 2a). We found the highest rates of wet deposition in northwestern of Mexico City. The rest of the valley lacks monitoring stations. The only existing record is from a natural protected area at the north of the Valley, whose wet deposition is always below 5 kg ha^−1^ year^−1^, ^17^.

**Figure 1.** Localization of the Valley of Mexico. Red and yellow dots represent the spatial distribution of the air quality network stations of the wet deposition and the automatic monitoring network for NOx, respectively. This network is located mainly in the Mexico City and its metropolitan area. Green triangles represent the sites where biomonitors were collected throughout the Valley. The red line delimits the basin, the white line indicates state division, and the blue line shows Mexico City limits. The map was created with ArcGIS 10 (Esri, Redlands, California, USA). Image data: Google Earth; image date: 5 September 2016.

**Figure 2.** Spatial distribution of the total wet deposition in Kg N ha^−1^ year^−1^ during 2014 (A), and atmospheric concentration of NOx in ppb (B). Data is available for public access in the website of the Mexico City government (http://www.aire.cdmx.gob.mx/default.php). The map was created with ArcGIS 10 (Esri, Redlands, California, USA). Image data: Google Earth; image date: 5 September 2016.

For NOx concentrations, the northern part of Mexico City is the zone with the highest concentrations reaching a cumulative of 78.8 ppb during the dry season and 40.6 ppb during the rainy season (Figure 2b). The lowest concentrations of NOx were found in the south and the east part of Mexico City which reached 21.8 ppb and 9.2 ppb during dry and rainy season respectively (Figure 2b).

### Nitrogen relations of biomonitors

#### Anaptychia sp

The nitrogen content of *Anaptychia sp*. ranged from 1.4% (dry mass) in a natural protected area at the north of the Valley to 5.0% in an urban site in Mexico City during the dry season (Figure 3a). We observed the same pattern during the rainy season, in which the nitrogen content ranged from 1.3 to 4.6%. The interaction between site and season was significant for all parameters measured (Table 1). A notable finding is that the highest nitrogen deposition of 50 kg N ha^−1^ year^−1^ did not produce the highest nitrogen content in this lichen, which was observed in a site whose nitrogen deposition reached 31.5 kg N ha^−1^ year^−1^.

**Table 1.** Two-way ANOVA for responses of potential biomonitoring organisms growing in the Valley of Mexico.

| | | d.f. | %N (dry weight) | | C:N ratio | | $\delta^{15}\text{N}$ (‰) | |
| --- | --- | --- | --- | --- | --- | --- | --- | --- |
|  |  |  | F | P | F | P | F | P |
| *Anaptychia* sp. | Site | 33 | 26.16 | <0.001 | 34.78 | <0.001 | 29.04 | <0.001 |
|  | Season | 1 | 7.83 | 0.005 | 10.14 | 0.002 | 0.99 | <0.001 |
| | Site × Season | 33 | 3.18 | <0.001 | 3.22 | <0.001 | 2.34 | <0.001 |
| *Grimmia* sp. | Site | 31 | 34.33 | <0.001 | 40.73 | <0.001 | 123.04 | <0.001 |
|  | Season | 1 | 1.47 | 0.226 | 30.02 | <0.001 | 2.95 | 0.087 |
| | Site × Season | 31 | 5.27 | <0.001 | 3.25 | <0.001 | 4.95 | <0.001 |
| *Fabronia* sp. | Site | 29 | 24.92 | <0.001 | 15.50 | <0.001 | 46.87 | <0.001 |
|  | Season | 1 | 2.57 | 0.11 | 6.14 | 0.014 | 1.44 | 0.232 |
| | Site × Season | 29 | 5.80 | <0.001 | 3.93 | <0.001 | 6.62 | <0.001 |
| *Tillandsia recurvata* | Site | 21 | 34.82 | <0.001 | 33.92 | <0.001 | 57.47 | <0.001 |
|  | Season | 1 | 96.05 | <0.001 | 6.53 | 0.011 | 136.29 | <0.001 |
| | Site × Season | 21 | 16.02 | <0.001 | 3.97 | <0.001 | 7.56 | <0.001 |

**Figure 3.** Spatial distribution of the nitrogen content (A, C, E, G) and δ^15^N values (B, D, F, H) for *Anaptychia* sp. (A, B), *Grimmia* sp. (C, D), *Fabronia* sp. (E, F) and *Tillandsia recurvata* (G, H). The map was created with ArcGIS 10 (Esri, Redlands, California, USA). Image data: Google Earth; image date: 5 September 2016.

The C:N ratio ranged from 7.5 to 39.1, increasing from the urban to the rural sites during the dry season; it ranged from 9.4 to 34.1 during the rainy season. Similarly, the δ^15^N values ranged between –9.4 and 5.2‰, being more positive in northeastern and downtown Mexico City and in southeastern Pachuca, contrasting with negative values from rural areas (Figure 3b). A linear regression showed no direct relationship between nitrogen deposition and the parameters measured for this lichen (R^2^=0.14 for nitrogen content, R^2^=0.07 for the C:N ratio and R^2^=0.16 for the δ^15^N).

#### Grimmia sp

We found significant differences between the nitrogen content of mosses from a natural protected area at the north of the valley that reached 1.3% where the deposition was 5.0 Kg N ha^−1^ year^−1^, with those of a site at the north of Mexico City of 3.8% where the deposition reached 40 Kg N ha^−1^ year^−1^ (Figure 3c). We observed no differences in the nitrogen between for the dry and rainy season (Table 1). A linear regression showed that the atmospheric concentration of NOx had no effect on the nitrogen content (R^2^ < 0.01). The C:N ratio was higher in rural areas and decreased significantly in the urban ones, it ranged between 9.5 and 33.6. On the contrary, NOx concentration had no effect on the C:N (R^2^ < 0.01).

We found the most negative δ^15^N values of –7.0‰ in a rural area and positive ones of 6.6‰ in urban sites where wet deposition was lower than 35 Kg N ha^−1^ year^−1^, exceeding this point δ^15^N became negative reaching –5.3‰, as occurs with mosses growing at 40 Kg N ha^−1^ year^−1^ (Figure 3d; Figure 4). We also observed positive values in the city of Pachuca at the north of the Valley. Season and NOx concentration showed no effect on the δ^15^N of this moss (Table 1).

**Figure 4.** Relationship between wet deposition of ammonium (green circles), nitrate (blue squares), and total deposition, NH_4_^+^ + NO_3_^−^ (red triangles) during 2014 and the δ^15^N values of the moss *Grimmia* sp.

#### Fabronia sp

The nitrogen content ranged between 1.7 and 3.4% (dry weight) across the Valley. We found no differences between the seasons, but significant differences between the rural areas and some sites in Mexico City (Table 1; Figure 3e). Wet deposition had no effect on the nitrogen content for the mosses from Mexico City. For example, it only increased by 0.3% when nitrogen deposition when from 24.1 to 48.3 Kg N ha^−1^ year^−1^. The site and the season were statistical significant (Table 1). The NOx concentration had no effect on the nitrogen content of this moss. In addition, wet deposition, nor the NOx concentration resulted in alterations on C:N ratio, which ranged from 9.0 and 29.5.

The δ^15^N ranged from –6.6 to 7.8‰. We found the most negative value in a semirural site at the north of the Mexico City (Figure 3f), while a δ^15^N of –6.0 ± 0.2‰ was observed at the south, where the deposition reached 37.6 Kg N ha^−1^ year^−1^. The most positive value was found in the eastern border of Mexico City, where the nitrogen deposition was 33.7 and 27.1 Kg N ha^−1^ year^−1^ during 2013 and 2014 respectively. The urban environment in the city of Pachuca also had a considerable effect on the isotopic composition of this moss, as positive values were the rule. Significant differences were found between rural areas and the urban ones. In contrast, the season showed no effect on the isotopic composition of this moss (Table 1). We found a weak relationship between the rate of nitrogen deposition and the δ^15^N values (R^2^ = 0.2; Figure 5).

**Figure 5.** Relationship between wet deposition, of ammonium (green circles), nitrate (blue squares) and total deposition, NH_4_^+^ + NO3 ^-^ (red triangles) during 2014 and the δ^15^N values of the moss *Fabronia* sp.

#### Tillandsia recurvata

The nitrogen content for *Tillandsia recurvata* ranged from 0.8 to 3.6% during the dry season and from 1.0 to 2.2% (dry weight) during wet season (Figure 3g). The highest content was observed in the northern part of Mexico City and was significantly different from the lowest value found in a semirural site in the central part of the Valley. At the same time, bromeliads collected in the city of Pachuca had a similar nitrogen content as those from semirural areas of the Valley (Figure 3g). However, this variation was not due to wet deposition. Instead, it responded positively to the NOx concentration (Figure 6a). For example, the nitrogen content reached 3.6% in an urban site where the NOx reached 57.4 ppb, contrasting with the lowest nitrogen content that reached 0.8% in a rural site where NOx presumably was lower to 5 ppb, based on the lowest recorded concentration in Mexico City. Wet deposition had no effect on the C:N ratio of this bromeliad, but it was strongly affected by the concentration of NOx during both seasons (Figure 6b). For example, we found the lowest C:N ratio of 15.9 where the NOx reached 50.3 ppb, and the highest C:N of 40 where NOx concentrations presumably were lower than 5 ppb. The δ^15^N values for *T. recurvata* were negative in rural areas and became positive in the cities, varying from −5.0 to 4.4‰ during the dry season and −7.7 to 5.1‰ during the wet season (Figure 3h). The δ^15^N values responded positively to the NOx concentration (R^2^=0.49; Figure 6c), while the wet deposition had no effect on δ^15^N values (R^2^=0.01). the data from two sites (Figure 6c open circles) were excluded from the analysis because topographic and exposure to pollutants were non-representative, clearly skewed the isotopic signature of negative values. In particular, one of the sites is a zoological park with large mammals that is crossed by a stream that was visible polluted. In turn, the second site were from a small ravine with dense vegetation, where the diffusion of NOx was probably reduced and where biogenic emissions were negative.

**Figure 6.** Relationship between NOx concentration during the 2014 dry season and the nitrogen content (A), C:N ratio (B), and the δ^15^N values (C) of the bromeliad *Tillandsia recurvata.* Open circles were excluded from the regression analysis because they were collected from non-typical environmental conditions that skewed the isotopic signatures of *T. recurvata* to very negative values.

## Discussion

The distribution of nitrogenous pollution and deposition in Mexico City generally matched that of the sources of emission. For instance, the highest values were measured in the northern part of the city, where the majority of the manufacturing industries are located along numerous important and busy motorways^18^. Indeed, motor vehicles, followed by industrial emissions, are the most important source of NOx in México City, whose concentration is directly measured by the city government from the air^18,19^. In addition, nitrogenous emissions from the agricultural zone between Mexico City and Pachuca substantially contribute to the observed pattern, given predominant north-to-south winds^18^. In this case, NHx originated from both agricultural and industrial activities are measured from wet deposition.

The nitrogen content of lichens has been utilized as an effective indicator of nitrogen deposition. For example, three species of lichens can record nitrogen inputs of wet deposition below 10 Kg N ha^−1^ year^−1^, point at which they saturate ^12,20^. We observed a threshold of nitrogen tolerance for lichens from the Valley, when the deposition reached 31.5 kg N ha^−1^ year^−1^ above which they could not take up additional nitrogen. We observed a similar pattern for the δ^15^N, while the negative values from rural areas were the result of low rates of deposition, in urban areas high rates of deposition resulted in positive isotopic values, however, above the threshold of saturation the isotopic values turned negative, as has been already observed in other species of lichen ^21^. Their isotopic composition responded more to wet than dry deposition as was observed for mosses. Additionally, different studies show a point of saturation when studying throughfall deposition ^20, 21^.

The nitrogen content of moss tissues is affected by the rate of deposition and responds to the distance to urban centers ^14,22-24^. We also observed an important influence of urban centers for the nitrogen content of mosses from the Valley. Their δ^15^N values are affected by the rates of deposition and the prevalent pollutant (e.g. wet NH_4_^+^ or NO_3_^−^; dry NHx or NOx,), being more commonly negative in mosses from rural areas ^25-28^. The observed negative δ^15^N for mosses from rural areas resulted from uptake of NH_4_^+^ derived from fertilizers and livestock emissions which have characteristic negative δ^15^N values ^27,29,30^. Mosses from urban areas of the Valley such as Pachuca, as well as some areas of Mexico City where the wet deposition was below 35 Kg N ha^−1^ year^−1^ had positive δ^15^N, suggesting that they more likely take up NO_3_^−^ derived from NO x of fossil fuel burning and industrial activities which typically display positive δ^15^N ^29^. Another possible mechanism influencing the isotopic composition of these mosses is the gas-particle conversion process, in which isotopic enrichment occurs for the aerosol nitrogen resulted, producing positive δ^15^N ^31^.

Mosses take up NH_4_^+^ preferentially over NO_3_^−^ because less energy is needed in its assimilation ^32^. Additionally, high deposition rates can cause the inhibition of nitrate reductase, reducing NO_3_ ^-^ assimilation ^27,33,34^. This occurs to some degree when the deposition reaches 10 Kg N ha^−1^ year^−1^, but when it exceeds 30 Kg N ha^−1^ year^−1^ the nitrate reductase become completely inhibited not only precluding for the NO3 ^−^ be uptake leading to its loss by leaching ^27,35,36^. Nitrate reductase inhibition also has an important effect on the isotopic composition of mosses because it favors the take up of NH_4_^+^ that typically has negative δ^15^N values ^30^. In the Valley of Mexico, mosses δ^15^N became negative when the rates of wet deposition exceeded 35 Kg N ha^−1^ year^−1^, which suggests that nitrate reductase was indeed inhibited, allowing NH_4_^+^ uptake, a pollutant that represented 35% of the total wet deposition in Mexico City during 2014 ^18,36,37^. The atmospheric concentration of Nox had no effect on the nitrogen content nor the isotopic composition of either moss considered here because when NH_3_ is the prevalent nitrogenous pollutant in the atmosphere, the nitrogen content of the mosses can increase more than from wet deposition. However, in Mexico City the prevalent nitrogenous gas pollutant is NOx, suggesting that nitrogen content and the δ^15^N of the mosses responded more to wet deposition than gaseous pollutants ^27,38^.

Despite the high rates of wet deposition recorded in Mexico City which exceeded 50 Kg N ha^−1^ year^−1^ in some areas, neither the nitrogen content nor the C:N ratio of *Tillandsia recurvata* were directly affected. This occurs because raindrops cannot be absorbed by the non-absorptive roots of these plants ^39^. Instead, the nitrogen was taken up as NOx by the leaves of this bromeliad because it can absorb particles and gasses from the air thanks to stomatal gas exchange and the trichomes present in the surface ^39^. This is evident from the close relationship found here between the nitrogen content and the NOx concentration in the Valley. This is similar to what occurs for other atmospheric bromeliads, whose nitrogen content is higher in the vicinity of highways than further away ^40-42^. The seasonal differences found for the nitrogen content appeared to respond to the phenology of this bromeliad which grows after the rainy season. Indeed, the NOx were dragged from the atmosphere to the ground surface during rainy season reducing its biological availability^18,43^.

Both the nitrogen content and the isotopic composition of *T. recurvata* were determined by the predominant anthropogenic activity of the sites where the plant was collected. Bromeliads take up NOx with positive δ^15^N from industrialized and densely populated areas, contrasting with the negative emissions originated from biogenic sources, the soil and livestock waste ^44,45^. The positive δ^15^N found for *T. recurvata* suggests no nitrate reductase inhibition, but it is likely that NOx concentrations higher than 80 ppb may result in some inhibition of this enzyme. This could be one of the reasons why *T. recurvata* was not found in sites with concentrations of NOx higher to 80 ppb.

The distinct responses observed for each biomonitor evaluated here are a result of the ecophysiological traits of each functional type. Indeed, the lack of a cuticle and a vascular system for mosses, allows a rapid/ready assimilation of incoming pollution, mainly from wet deposition. However, their weak response to NOx levels could be an effect of plant phenology, as the mosses remain dormant during the dry season, so the uptake of the gas is not physiologically possible. For the case of the tillandsias, the fact that they are adapted to a water and nutrient-limiting environment was the cause for its suitability for monitoring NOx, as all their nutrition is atmospheric and their water is intercepted from the atmosphere ^46^. In addition, their potential uptake of wet deposition is inherently limited by their small area to volume ratio that in turn reduces water loss ^47^.

Also, their rates for taking up water are low, so exposure to more benign environments does not lead to increased water uptake as we have found experimentally ^48^. Of course, caution must be taken when utilizing a biomonitoring method utilizing spontaneously occurring plants, as their nitrogen parameters can also respond to various factors other than nitrogenous pollution. For instance, if dewfall is substantial during the dry season, mosses can remain physiologically active, and thus be able to assimilate NOx. Also, water availability and extreme temperatures can have an effect on the N relations of plants. For instance, high daytime temperatures and very low relative humidity during the dry season or a very cloudy sky during the rainy season can both lead to stomatal closure during the night for the CAM bromeliads. In either case, uptake of NOx will not occur. Other factors to consider are pathogens and herbivores ^49^. For these reasons, specific biomonitors need to be developed for each region of interest, taking in consideration the particular environmental conditions and the ecophysiological traits of potential biomonitors ^11^. For the case of the Mexico valley, and by possible extension to the Trans-Mexican Volcanic Belt, *T. recurvata* can be utilized for monitoring NOx along with a moss that records wet deposition. This approach can become a useful tool for determining the status of nitrogenous pollution in regions where air quality monitoring networks are not available. Especially in mid-sized cities and surrounding areas where the saturation thresholds of these organisms has not been reached. Finally, the utilization in tandem of these organisms can inform an early alert for avoiding health problems for both ecosystems and humans.

## Methods

### Study area

The Valley of Mexico is located in central Mexico spanning 7500 km^2^, with an average elevation of 2240 m ^50^. Annual precipitation ranges from 600 mm at the center of the valley to 1300 mm in the surrounding mountains. The predominant winds blow from the northeast and northwest ^51^. Mexico City sits at the southern edge of the Valley with a population of 20 million. At the northern edge, Pachuca, the capital of the state of Hidalgo, has a population of 3 million. Additional, settlements interspersed in the valley with industrial or agricultural activities comprise the rest of the 30 million inhabitants of this basin ^10^.

### Nitrogen environment

The Mexico City environmental authority has deployed an air quality network of 16 monitoring stations for wet deposition (Fig. 1). This network collects data during the rainy season from May to November. The nitrogen collected consists of dissolved NO3 ^-^ and NH_4_^+^, so that the total nitrogen is the sum of both forms of deposition (NO_3_^−^ + NH_4_^+^). This is the dissolved inorganic nitrogen (DIN), biologically available ^27^. For this study, the rates of wet deposition measured by this monitoring network during 2014 were utilized for the analyses described below. This air quality network also has 27 monitoring stations for measuring the atmospheric NOx concentration (NO + NO_2_) year-round, 24 hours a day. We utilized the mean seasonal concentration in ppb during dry season (November 2013 to April 2014) and during the rainy season (from May 2014 to October 2014). Data were obtained from the website of the environmental authority of the Mexico City (http://www.aire.cdmx.gob.mx/default.php).

### Biomonitoring

We selected four species of three types of atmospheric organisms, the lichen *Anaptychia* sp., the mosses *Grimmia* sp. and *Fabronia* sp., and the bromeliad *Tillandsia recurvata* (L). These organisms have been utilized as monitors of nitrogen deposition and are abundant in the Valley of Mexico ^14,15,21^. We collected tissue samples from 5 individuals of each species growing on different substrates from 36 sites throughout the valley, including urban parks, agricultural sites, and natural protected areas, for the dry season on (8-20 May) and, the wet season 3-15 November of 2014. The distribution of the sites within the basin was determined by the occurrence of the biomonitoring species and by the complex nature of the landscape, which precluded the collection of samples from a regular grid ^52^.

The tissue samples were dried at 60°C in a gravity convection oven until reaching constant weight. The dried tissues were ground to a fine powder in a ball mill (Retsch MM300; Retsch, Vienna, Austria), wrapped into tin capsules (Costech Analytical, Inc. Valencia, California, USA), and weighed with a microbalance (0.01 mg, Sartorius, Göttingen, Germany). For each sample, both the nitrogen content and their isotopic signature were determined at the Stable Isotope Facility, University of Wyoming (Laramie, Wyoming, USA), with a Carlo Erba EA 1110 elemental analyzer (Costech Analytical Inc., Valencia, CA, USA) attached to a continuous flow isotope ratio mass spectrometer (Finnigan Delta Plus XP, Thermo Electron Corp, Waltham, MA). Nitrogen isotope ratios, reported in parts per thousand, were calculated relative to atmospheric air standards. The analytical precision for the δ^15^N was 0.3 ± 0.07‰ (SD). The natural abundances of ^15^N were calculated as:

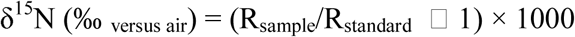

where, R is the ratio of ^15^N/^14^N for nitrogen isotope abundance for a given sample ^53,54^.

### Statistical analyses

We utilized linear regressions to determine the relationship between total wet nitrogen deposition and the nitrogen content (% dry weight), the C:N ratio as well as the δ^15^N of the organisms considered. The same was done for the concentration of atmospheric NOx. Data accomplished normality in both cases ^30^. We performed two way ANOVAs (factors were site and season) followed by a post hoc Holm–Sidak tests (*p* ≤ 0.05) to determine differences in the nitrogen content (% dry weight), the C:N ratio, as well as the δ^15^N values of the tissue samples. All statistical analyses were conducted with Sigmaplot 12 (Systat Software Inc. USA).

### Geostatistical analyses

We utilized the ordinary Kriging method (a geostatistical gridding tool for irregularly spaced data ^55^) to determine the rates of wet deposition, and the concentration of NOx in the sites where no monitoring station was available for the area covered by the monitoring network. Likewise, we modelled the geographical distribution of the nitrogen content, C:N ratio, and the δ^15^N values for the biomonitors, as well as the data from the monitoring network with ArcGIS 10 (Esri, Redlands, California, USA).

### Data availability

The datasets generated and/or analyzed during the current study are available from the corresponding author on reasonable request.

## Acknowledgments

We thank funding by the Dirección General de Asuntos del Personal Académico, (PAPIIT IN205616), and institutional funds of the Instituto de Investigaciones en Ecosistemas y Sustentabilidad, Universidad Nacional Autónoma de México. EADA held a generous graduate research fellowship from Consejo Nacional de Ciencia y TecnologÍa, México. We also thank the generousity of the personnel of Zona Arqueológica de Teotihuacán, especially to Ms. V Ortega.

## Contributions

Both authors planned the study, conducted field work, took part in the preparation of the manuscript, have reviewed the results, and have approved the final version of this manuscript. E.A.D.A. analyzed the data, prepared all of the figures, and wrote the initial version of the manuscript. E.d.l.B. procured funding, revised and approved the manuscript, and drove in the field and planned a gastronomic tour in the Mexico Valley.

